# High diversity and heterogeneity define microbial communities across an active municipal landfill

**DOI:** 10.1101/2021.03.26.437222

**Authors:** Alexandra H. Sauk, Laura A. Hug

## Abstract

Global waste production is increasing rapidly, with the majority of waste destined for landfills. Microbial communities in landfills transform waste and generate methane in an environment unique from other built and natural environments. Previous work has largely considered landfill microbial diversity only at the phylum level, identifying complex and variable communities. The extent of shared organismal diversity across landfills or over time and at more precise levels of classification remains unknown. We used 16S rRNA gene amplicon and metagenomic sequencing to examine the taxonomic and functional diversity of the microbial communities inhabiting a Southern Ontario landfill. The diversity of microbial populations in leachate and groundwater samples was correlated with geochemical conditions to determine drivers of microbial heterogeneity. Across the landfill, 25 bacterial and archaeal phyla were present at >1% relative abundance within at least one landfill sample. The *Patescibacteria*, *Bacteroidota*, *Firmicutes*, and *Proteobacteria* had the highest relative abundances, with most other phyla present at low (<5%) abundance. Below the phylum level, very few populations were identified at multiple sites, with only 121 of 8,030 populations present at five or more sites. This indicates that, although phylum-level signatures are conserved, individual landfill microbial populations vary widely. Significant differences in geochemistry occurred across the leachate and groundwater wells sampled, with calcium, iron, magnesium, boron, meta and para xylenes, ortho xylenes, and ethylbenzene concentrations contributing most strongly to observed site differences. This study illustrates that leachate microbial communities are much more complex and diverse within landfills than previously reported, with implications for waste management best practices.

## Background

The environmental impact and monetary cost of municipal solid waste (MSW) storage and management are growing concerns for municipalities and countries around the world. MSW includes food waste, paper, plastics, and metals [1]. MSW generation has increased exponentially with rising populations, increased development, and urbanization [2,3]. By 2025, the global annual production of waste will reach an estimated 2.2 billion tons, and is not predicted to hit a maximum in this century, reaching 4 billion tons annually in 2100 [1,2]. This is an unsustainable rate of increase.

Landfills are the most common end point for MSW in many countries, including Canada, the United States, and China. Landfills are the third largest contributor to anthropogenic methane emissions, contributing 11% of annual global methane emissions and making them a focus area for mitigating climate change [1,4,5]. Landfill sites for MSW range from open dumping to sanitary landfills [3,6–9]. These disposal options can have varying levels of environmental protection and waste containment, and can be engineered to promote or prevent aerobic or anaerobic microbial metabolisms [3]. Waste degradation in landfills is controlled by the microbial communities within the landfill and the built characteristics of the landfill, such as leachate collection systems and cover soils [4,10]. The first three steps of the waste decomposition process are reliant on bacteria: hydrolysis; acidogenesis, including both fermentation and beta oxidation; and acetogenesis [10]. The last step, methanogenesis, is dependent on methanogenic archaea [10]. Landfill deposits are diverse, both chemically and physically, which can inhibit or prevent these microbial degradation processes [11]. Understanding the ecology and diversity of the bacterial and archaeal community structure in landfills will strengthen our understanding of the decomposition process, allowing for better control of methane production and more efficient waste management and contaminant mitigation strategies.

Despite recent interest in landfill microbial diversity [6–8,11–13], much is still unknown about microbial communities and their associated functions. Most early research focused on specific aspects of waste degradation in landfills and the microbes responsible, with particular interest in methane cycling [14,15] and cellulose degradation [16,17]. With the advent of high-throughput sequencing techniques like 16S rRNA gene amplicon sequencing, overall landfill microbial diversity and community composition have also been examined [11,12]. The most abundant phyla have been consistent across studies and include the *Firmicutes, Bacteroidota, Campylobacterota* (formerly *Epsilonproteobacteria* [18]) and *Proteobacteria* [8,10–12]. A number of rare and/or unclassified microorganisms have also been found in recent landfill studies, with some at high abundance [10,11]. These studies allow for functions to be inferred for microorganisms with well-characterized relatives, but the 16S rRNA gene cannot be used to infer functions for unclassified microorganisms that are uncultivated or newly described [10,12].

Several environmental or geochemical factors that influence microbial community composition and heterogeneity have been identified. Both landfill age and the age of waste has been correlated with microbial community composition characteristics [10–12,19]. These findings suggest that landfill microbial community composition changes over time, but more long-term studies are needed to determine if the change in community structure is predictable or if it is landfill-specific. Community composition was also correlated with moisture [8,12] and ammonium concentration [8,10]. Other chemicals that showed a link to microbial community composition included barium, chloride, sulfate, and copper [8,11]. Other chemical factors seem to affect microbial communities in a site-specific manner, and their effects will depend on the types of waste deposited and other geochemical conditions at each site of interest [11,19,20].

The study site for this research is a municipal waste landfill in southern Ontario, Canada that opened in 1972. This landfill is a conventional sanitary landfill with onsite waste sorting, compacting, and daily soil covers. The landfill is well-instrumented, with over 100 leachate wells (LW) across the site as well as three composite leachate cisterns (CLC). The leachate wells are routinely sampled by regional waste management staff to determine the chemical composition of the leachate. There are also groundwater wells (GW) bordering the landfill for monitoring the conditions of the adjacent aquifer and any leachate leaks (*e.g.*, there is on-going leachate infiltration from the area near LW3 into the aquifer near GW1, Figure 1). In order to understand waste degradation processes and the transformation and movement of contaminants within the site, it is important to understand how the microbial communities differ across the landfill. Here we combine metagenomic and 16S rRNA gene amplicon sequencing techniques to characterize the distribution, heterogeneity, and diversity of the microbial communities in a Southern Ontario municipal landfill. We additionally investigate how the observed microbial heterogeneity connects with geochemical conditions across the site.

**Figure 1:**
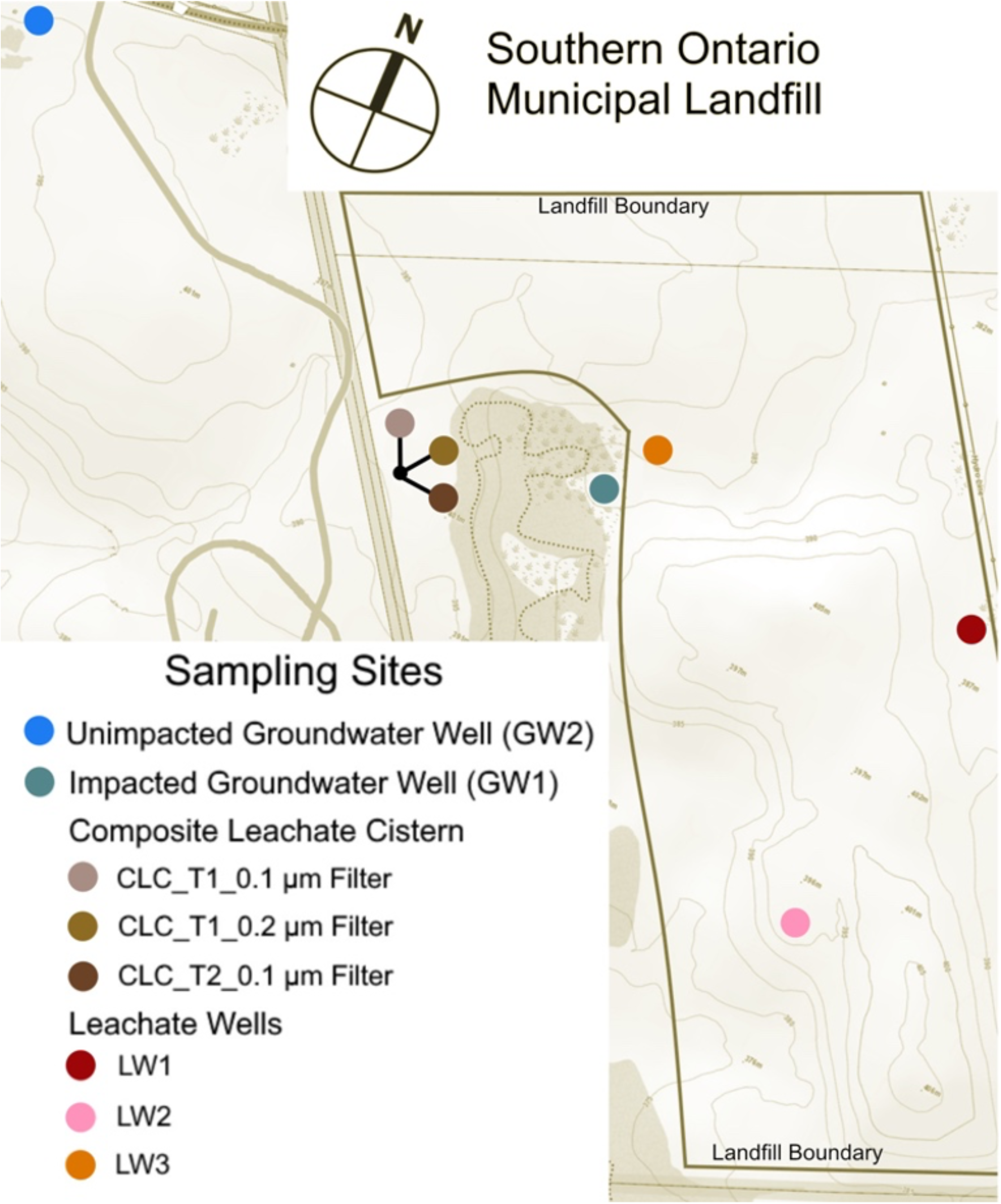
Map of landfill sampling locations at the Southern Ontario landfill. Two groundwater wells accessing the adjacent aquifer were sampled. The three samples from the leachate collecting cistern were all sampled from the same cistern at two time points. Two filter sizes were used for collecting microbial biomass on July 14, 2016 (CLC_T1_0.1 μm filter and CLC_T1_0.2 μm filter) and one filter size was used on July 20, 2016 (CLC_T2_0.1 μm filter). The three leachate wells are located within the active landfill, and leachate from these wells is pumped to the leachate collecting cistern. The catchment area for LW3 has an active leak infiltrating into the groundwater near GW1. GW1, the impacted groundwater well, shows geochemical evidence of leachate infiltration into the aquifer, where GW2 is upstream and shows no leachate chemical signature in the groundwater. The topographic map was modified from maps provided by the Ontario Ministry of Natural Resources and Forestry.

## Results

### Sampling and sequencing

Samples were collected from three leachate wells (LW1, LW2, and LW3), two samples from a composite leachate cistern at time points separated by one week (CLC_T1 and CLC_T2), and samples from two groundwater wells (GW1 and GW2) adjacent to the landfill (Figure 1). All biomass was harvested by serial filtration through a 3 μm glass fiber filter onto polyethersulfone 0.1 μm filters with the exception of the CLC_T1 sample, which was serially filtered through a μm and 0.1 μm filter. Total community DNA was extracted (see Methods for details). DNA from all seven filters, including two from CLC_T1 for the 0.1 μm and 0.2 μm filters, were processed by the US Department of Energy’s Joint Genome Institute (JGI) for 16S rRNA gene amplicon sequencing. Six DNA samples were sent to the JGI for metagenomic sequencing, assembly, and annotation: LW1, LW2, LW3, CLC_T1 (0.2 μm filter), CLC_T2, and GW1.

### Phylum level diversity

The 16S rRNA amplicon sequences were taxonomically classified and relative abundances were determined using QIIME2 [21]. From the 16S rRNA gene analysis, 8,030 exact sequence variants (ESVs) were identified across the sampled sites with an average of 1,147 ESVs per site. In tandem, metagenomic scaffolds were identified to the phylum level via placement on phylogenetic trees inferred based on the 16S rRNA gene and a suite of sixteen concatenated ribosomal proteins. Phylogenetic trees included 1,265 ribosomal protein marker-gene-encoding scaffolds and 2,306 metagenome-derived 16S rRNA genes for the ribosomal protein and 16S rRNA gene trees, respectively. The total number of medium or higher quality metagenome assembled genomes (MAGs) resolved from the metagenomes was 503. Taxonomy information was combined with scaffold coverage data to determine the relative abundances of phyla present in the landfill metagenomes. Twenty-five phyla were present at greater than 1% relative abundance in at least one landfill sample (Figure 2). Phylum level profiles were relatively consistent between the 16S rRNA gene amplicon and metagenomic sequencing data (Figure 2). A notable exception was the *Patescibacteria* (Candidate Phylum Radiation), which make up a comparatively reduced proportion of the 16S rRNA gene amplicon results (max relative abundance of 30.78%, in GW1) but exhibit the highest relative abundances in the metagenomic data (mean relative abundance of 34% and max relative abundance of 79%, in GW1, based on the coverage of the ribosomal protein-encoding scaffolds). The *Bacteroidota* (mean: 16%, max: 31.89% in CLC_T1), *Firmicutes* (mean: 10.19%, max: 28.74% in CLC_T1), and *Proteobacteria* (mean: 10%, max: 28% in LW2) were also highly abundant across the landfill sites.

**Figure 2:**
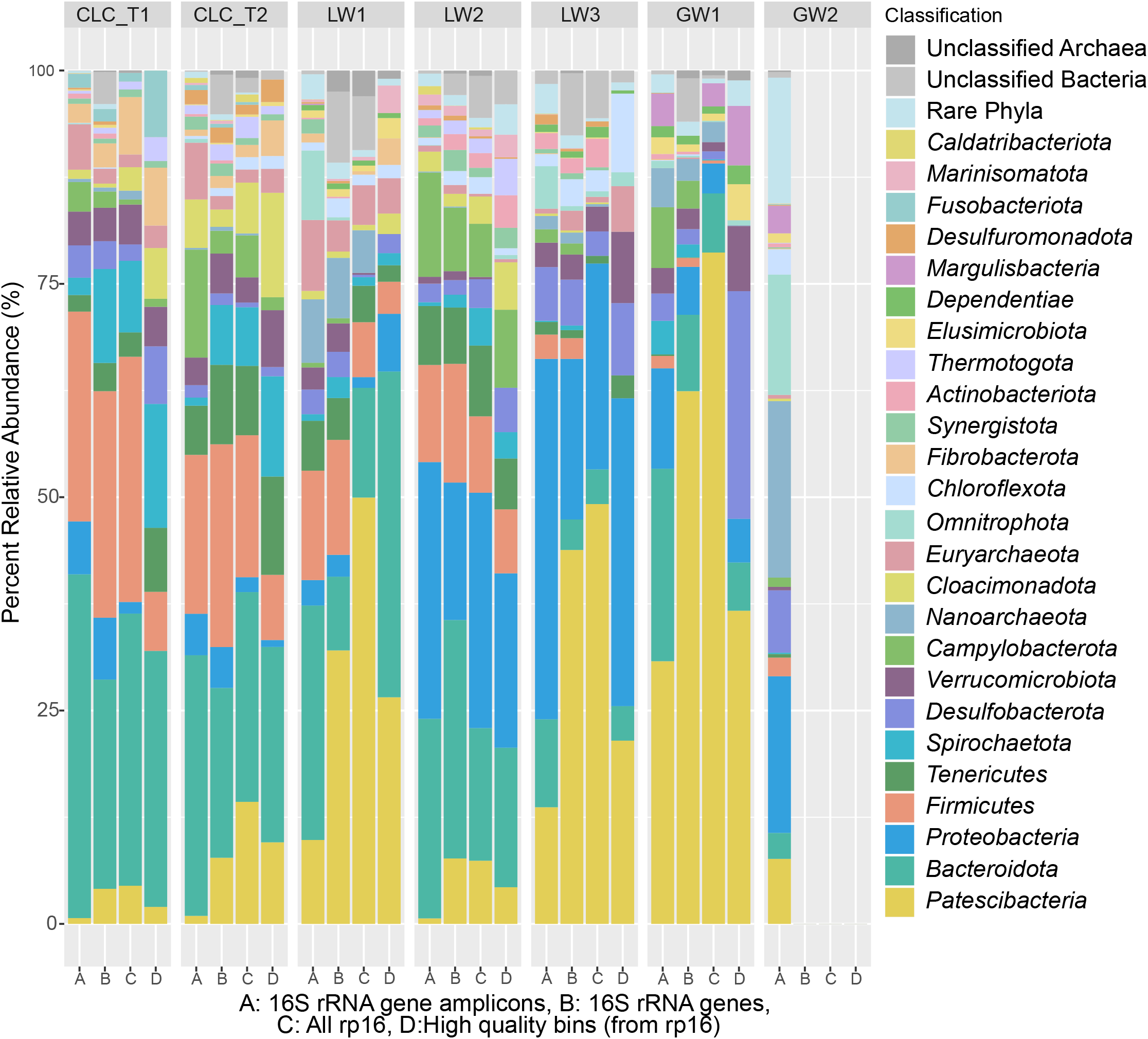
Relative abundances for phyla present at greater than 1% abundance in at least one sample. A) 16S rRNA gene amplicons; B) 16S rRNA genes derived from the assembled metagenomes; C) concatenation of 16 syntenic ribosomal proteins derived from the assembled metagenomes, with scaffold coverage as a proxy for abundance; and D) high quality bins (containing the 16 concatenated ribosomal proteins, and with abundances calculated from the ribosomal protein-encoding scaffolds’ coverages). The Composite Leachate Cistern at timepoint 1 (CLC_T1) is represented by the 0.2 μm filter’s sequence information, as the 0.1 μm filter showed highly similar results. Site GW2, the unimpacted groundwater well, did not yield sufficient biomass for metagenomic sequencing, thus only 16S rRNA gene amplicons are reported here.

### Alpha and beta diversity metrics

Alpha and beta diversity metrics were calculated based on the 16S rRNA gene amplicon sequences using QIIME2 and the phyloseq package in R [22]. All of the landfill samples had a Shannon index above 5.0 for the 16S rRNA gene amplicon data (Figure 3). There was no significant difference between the sample types (groundwater, leachate wells, leachate cisterns) when considering Faith’s phylogenetic diversity (Supplemental Figure 1A). The eight samples also exhibited high Pielou’s evenness (>0.74) with no significant differences between sample types (Supplemental Figure 1B).

**Figure 3:**
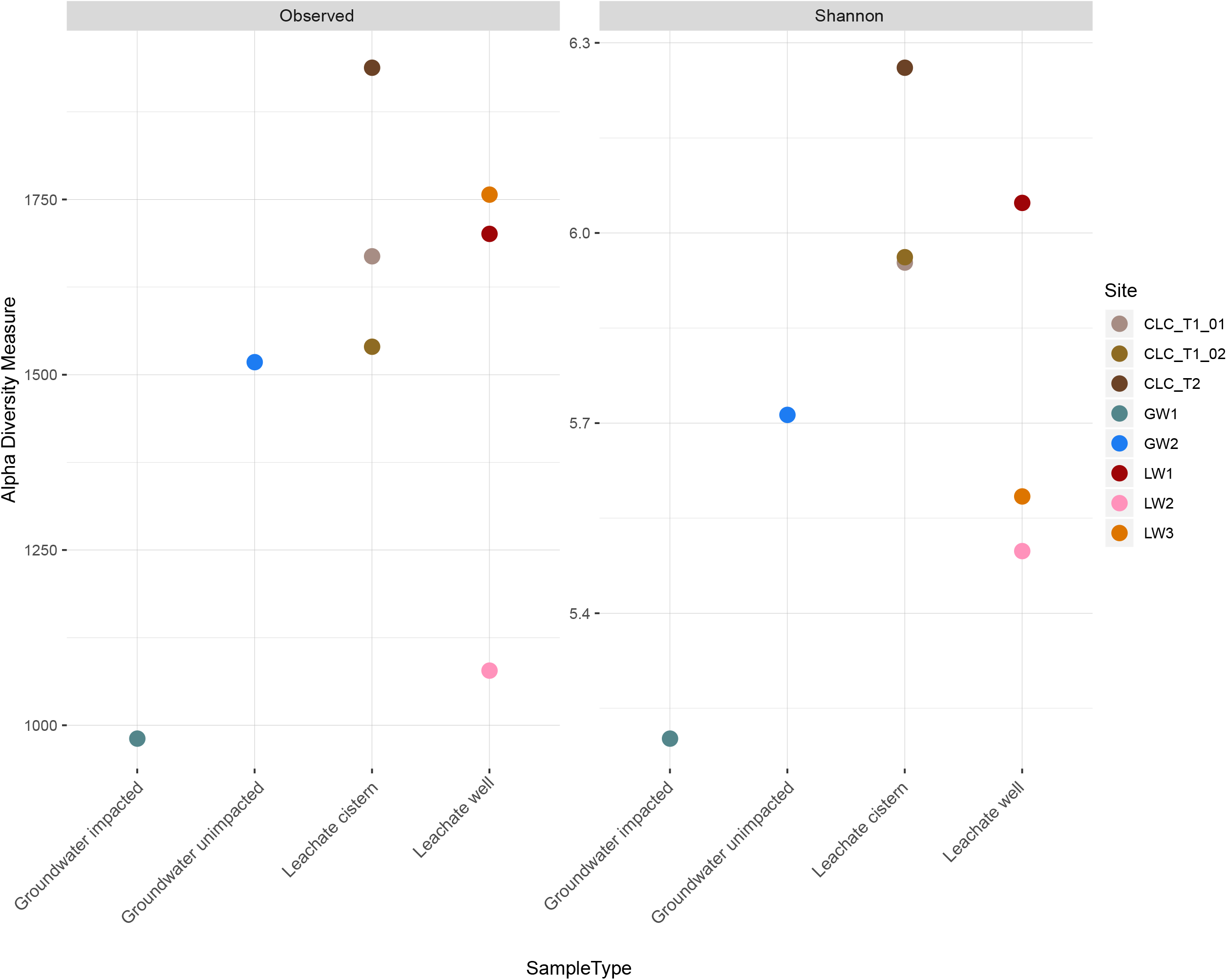
Observed diversity and Shannon index for the eight landfill-associated samples. Observed 16S rRNA gene amplicon ESVs fall between 986-1998 total ESVs for each sample, indicating similar levels of microbial community heterogeneity across sites. All sites show a high level of diversity, with Shannon indices greater than 5. Samples are grouped by sample type on the x axis.

Principle coordinates analysis (PCoA) plots using weighted and unweighted UniFrac distances based on 16S rRNA gene amplicon ESVs showed separation of the samples by type (Figure 4). The inclusion of abundance data in the weighted UniFrac analysis increased the explained variation on axes 1 and 2 by a combined 24.1%, suggesting that presence/absence and phylogenetic distance data implemented in the unweighted UniFrac are not sufficient to resolve the differences in beta diversity between sites in two dimensions. The inclusion of differences in abundance and overlap of ESVs between sites increased separation of the samples by type.

**Figure 4:**
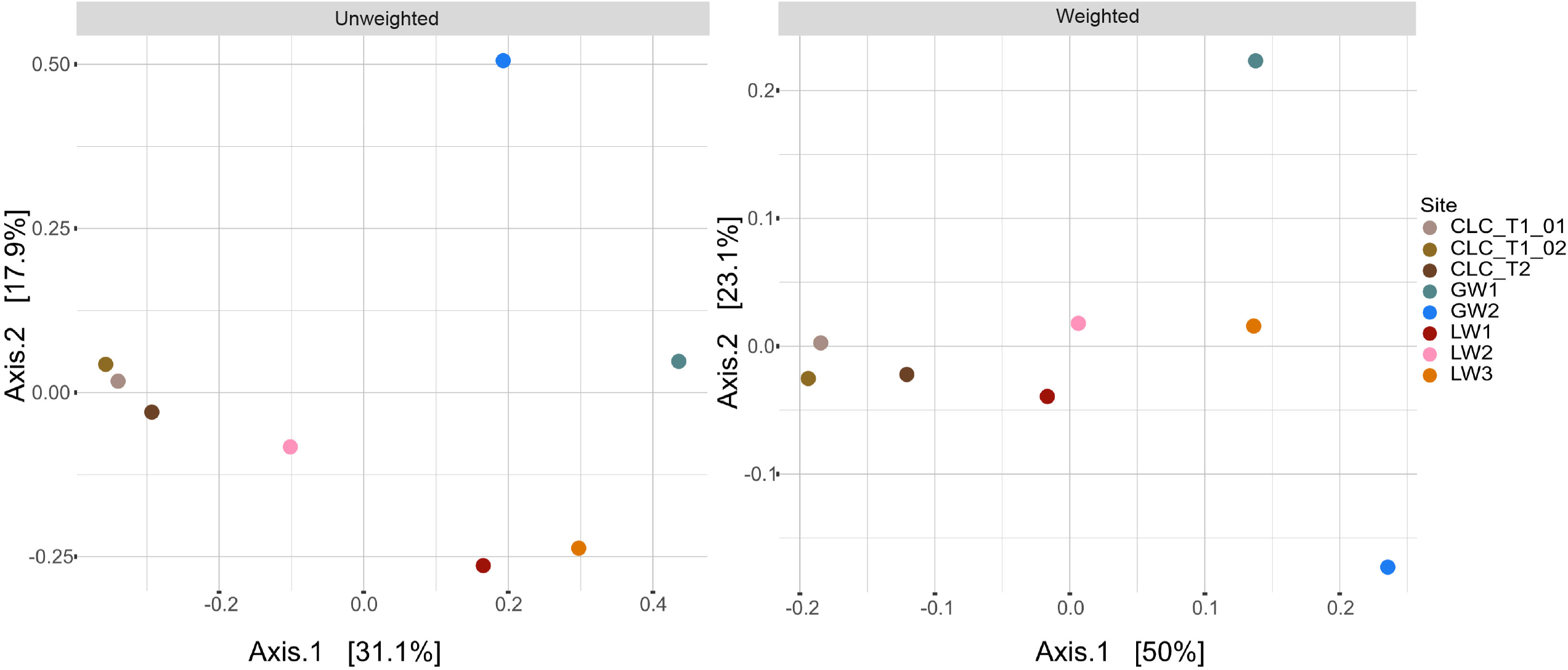
PCoA using unweighted and weighted UniFrac distances based on 16S rRNA amplicon ESVs for all sites.

### Diversity of exact sequence variants

The prevalence of 16S rRNA gene amplicon exact sequence variants (ESVs) was determined using phyloseq and visualized in ggplot2 [23] in R (Figure 5). The abundance of ESVs present in 5 or more sites were visualized using ggplot2 in R by phylum (Supplemental Figure 2). Although phylum level diversity was relatively consistent across the composite leachate cistern, leachate wells, and GW1 sample, the diversity at the ESV level is nearly entirely non-overlapping. The majority of ESVs identified from the top 25 phyla are present in only a single sample (Figure 5) with only 121 of 8,030 ESVs present across five or more samples (Figure 5 and Supplemental Figure 3). In addition to the top 25 phyla, ESVs belonging to LCP-89, *Micrarchaeota*, and an unclassified group of *Deltaproteobacteria* were also present in five or more sites. The abundance of phyla with populations across 5 or more phyla ranges by several orders of magnitude from 134 total ESV counts for *Elusimicrobiota* to 83,545 total ESV counts for *Bacteroidota* (Supplemental Figure 1). Of the 8,030 ESVs, 73.82% were found in only one sample and the number of ESVs shared between any two sites is at maximum 1,165 ESVs (Figure 5 and Supplemental Table 1). Principle component analysis (PCA) for all ESVs showed separation of composite leachate cisterns, leachate wells, and groundwater wells is driven by highly abundant ESVs (Figure 6A). When considering only ESVs present at five or more sites, LW2 is separated from LW1 and LW3 along PC2 and GW1 is separated from all other sites along PC1 (Figure 6B).

**Figure 5:**
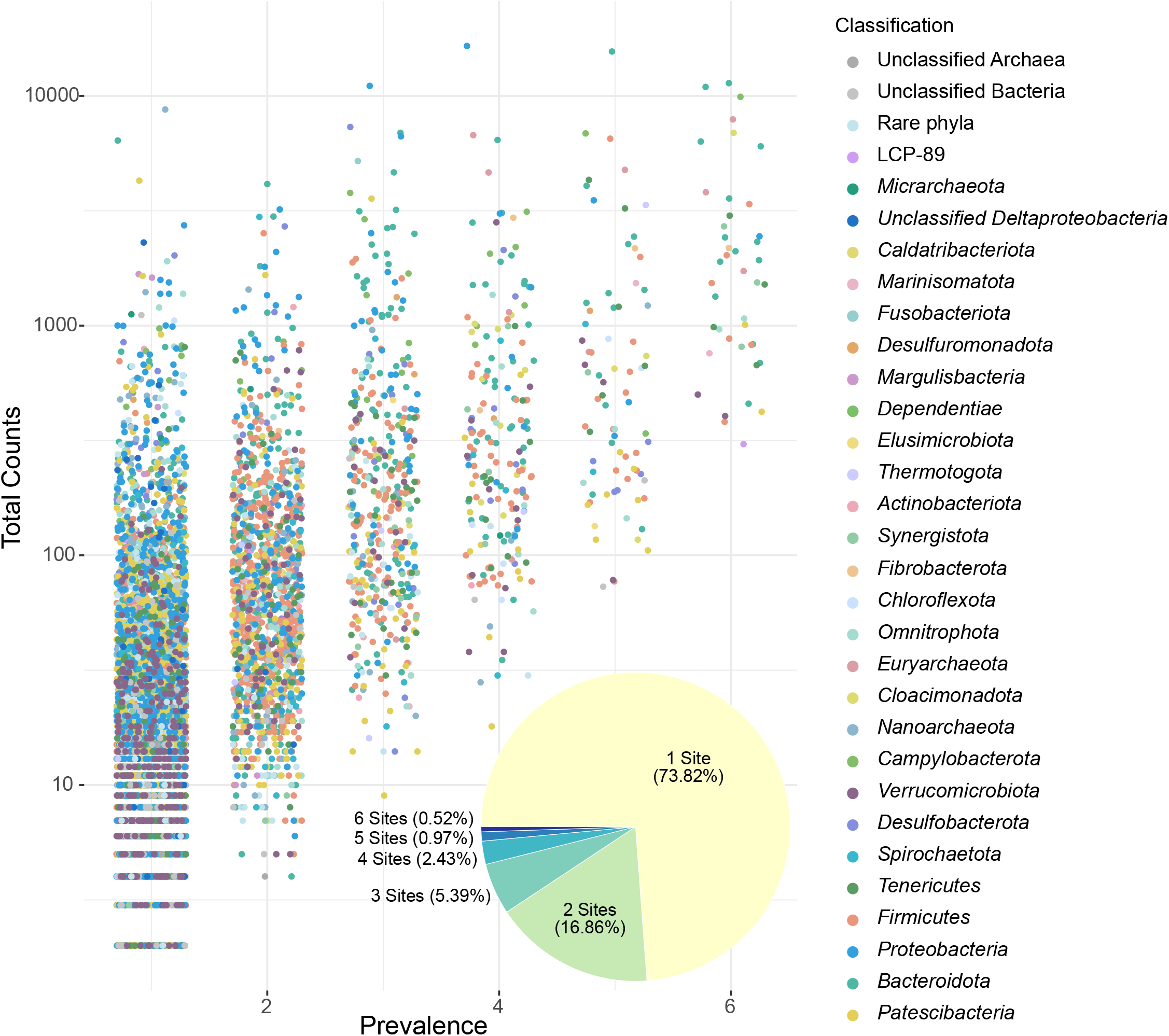
Prevalence of 16S rRNA amplicon ESVs across sampling sites. ESVs with individual counts greater than two from the top 25 most abundant phyla or present in four or more sites were included. The pie chart insert shoes the proportion of ESVs present in one to six sites. Data from CLC_T1 0.1 and 0.2 μm filters were combined, because these samples represent the same biomass from the same site and sampling time. The majority of ESVs occur in only one sample.

**Figure 6:**
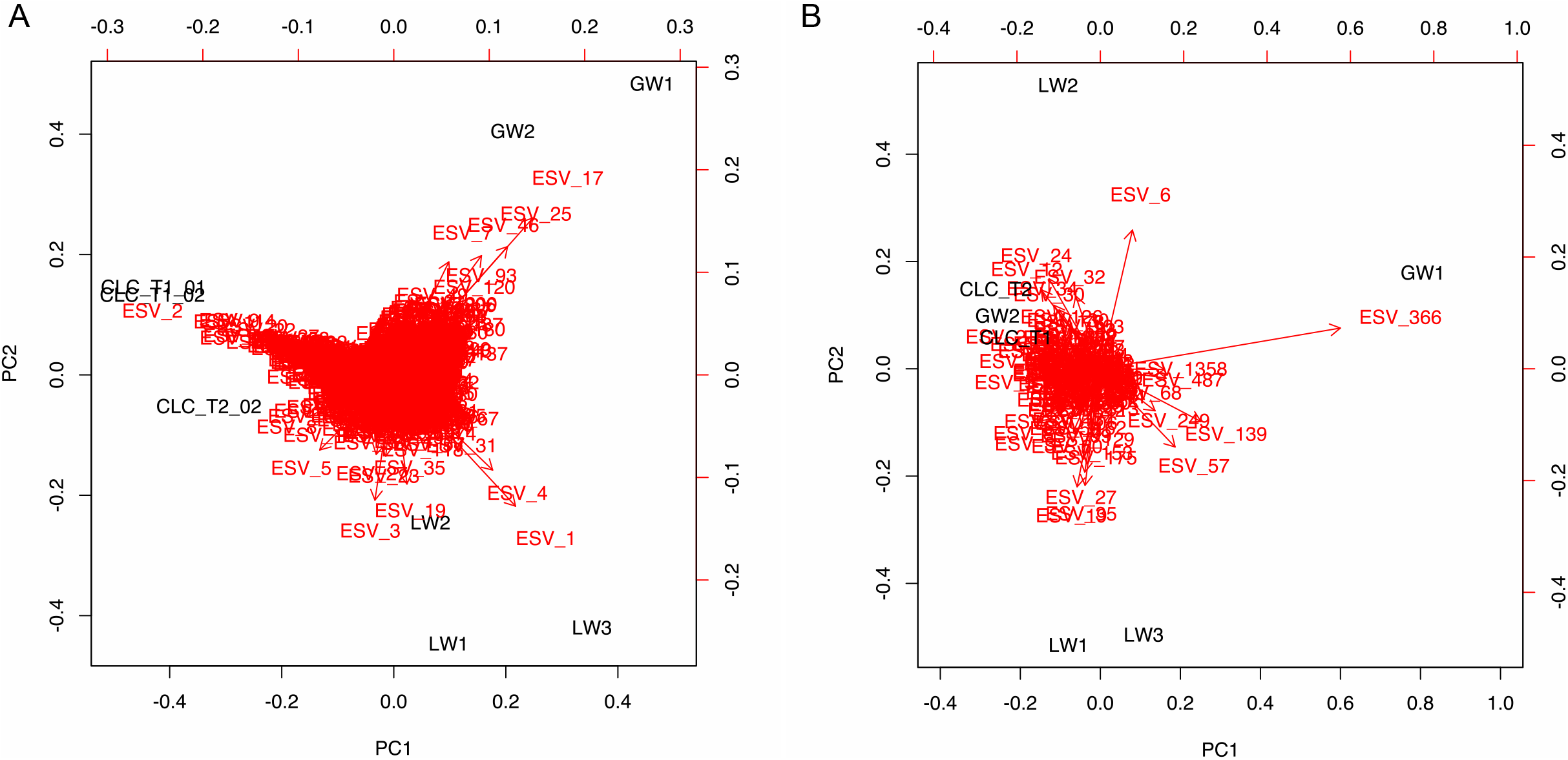
Principal component analysis for (A) all 16S rRNA gene amplicon ESVs and (B) those present at five or more sites. ESV count data was Hellinger transformed.

### Analysis of geochemical parameters

Geochemical parameters, including concentrations of volatile and non-volatile compounds, are measured quarterly by a contracted monitoring company. The statistical power available for analysis of geochemical parameters in the landfill was limited by the availability of data. Non-volatile compound measurements were only available for four sites and volatile compound measurements for five sites (Supplementary Table 2). Non-volatile and volatile compound concentrations varied significantly between sites when compared using an ANOVA (p< 9.14e−^14^ and p<2e^−16^, respectively), with a large range between sites for several non-volatile and volatile compounds (Figure 7). The date of measurement was not significant for either volatiles or non-volatiles when compared using an ANOVA (p=0.56 and p=0.73, respectively). The April and October 2016 measurements for the PCA analysis were averaged to estimate conditions during the July sampling for microbial biomass. Sodium and potassium were removed as outliers because their excessively high concentrations in LW2 (Supplementary Table 2) caused their variance to mask any differences in other compounds in the analyses. From the PCA, calcium, iron, magnesium, and to a lesser degree, boron contributed to the differences between the leachate wells and GW2 (Figure 7C). For the volatile compounds, nearly all of the observed variation is explained by PC1 (97.4%), largely due to the punctuated presence of m-& p- xylenes in LW1 and LW3, and of o. xylenes and ethylbenzene in LW1 (Figures 7B and 7D).

**Figure 7:**
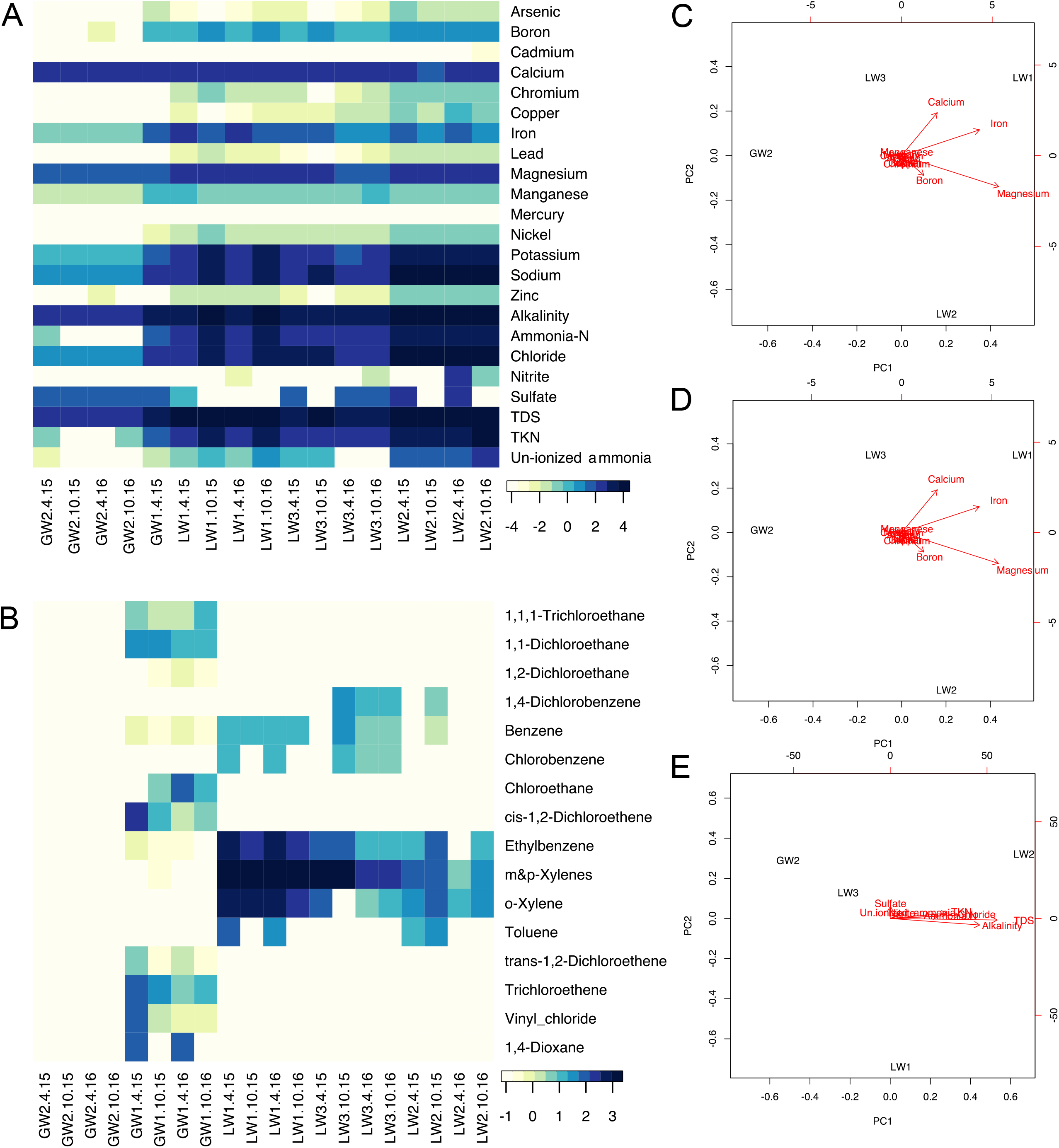
Environmental variation between landfill and aquifer sites for non-volatile (mg/L) and volatile (μg/L) compounds. A) Heat map of non-volatile compound concentrations and other site parameters (log10 transformed) across four dates for the three leachate wells and the unimpacted groundwater well. Only one date (April 2011) was available for the impacted groundwater well (GW1). Stars indicate low levels of cadmium and mercury not visible on the heatmap that may be biologically relevant. B) Heat map of volatile compound concentrations (log10 transformed) across four dates for the leachate and groundwater wells. C) Principal component analysis of metal concentrations at the three leachate wells and the unimpacted groundwater well. Metal concentration data was square root transformed. Sodium and potassium concentrations were much higher in LW2, so these compounds were removed to allow visualization of separation by other variables. PC1 explains 79.6% of variation and PC2 explains 18.4% of variation. D) Principal component analysis of volatile compound concentrations at the three leachate wells and the unimpacted groundwater well. Volatile compound concentration data were square root transformed. PC1 explains 97.4% of variation and PC2 explains 2.4% of variation. E) Principal component analysis for non-metal, non-volatile compounds, derived from measurements at the three leachate wells and the unimpacted groundwater well. Concentration data was square root transformed. PC1 explains 99% of the variation.

### Microbial diversity at groundwater wells

There are marked differences in the groundwater microbial communities from GW1 and GW2, the leachate-impacted and unimpacted wells, respectively. GW1 has a high abundance of Patescibacteria while also sharing a more similar phylum-level profile to the leachate wells than to GW2 (Figure 2). The sample from GW2 had insufficient microbial biomass for metagenomic sequencing, but 16S rRNA gene amplicon sequencing showed that GW2 has a distinct microbial community compared to all other sites, including a higher relative abundance of *Nanoarchaeaota* (20.7%) and *Omnitrophota* (14.1%) (Figure 2). The difference in microbial community composition between GW1 and GW2 is also reflected in their alpha diversity metrics. GW1 has the lowest Shannon index of the eight samples (Figure 3) as well as the lowest Faith’s phylogenetic diversity (Supplemental Figure 1A). A lower richness and evenness are expected in GW1, as the mixing of leachate and groundwater creates a suboptimal environment for microorganisms adapted to either environment [9].

## Discussion

### Phylum level diversity

The phylum level diversity in this landfill is generally consistent with other studies, where *Bacteroidota*, *Firmicutes*, and *Proteobacteria* are frequently detected as the most abundant bacterial phyla in landfills [8,11,12]. Interestingly, our study uncovered a high abundance of *Patescibacteria* in the landfill that had not been found in previous studies [11,12]. Previous landfill microbial diversity studies have relied on 16S rRNA gene amplicon sequencing, which may have systematically underestimated the abundance of *Patescibacteria*. Members of the *Patescibacteria* are often underrepresented in 16S rRNA gene amplicon studies due to primer mismatches and long insertions in the gene [24], but can be more robustly identified using metagenomic techniques [25].

### Alpha diversity

Shannon Indices above 5 for each site indicates that the landfill sites each have relatively high levels of microbial richness and evenness. This is greater than seen in some soil microbial communities, such as those in the Canadian prairies (1<*H*<4.5), and similar to others, such as in forest and agricultural land in Burgundy, France (4.5<*H*<6.1) [26,27]. Soil microbial communities show variation in richness and evenness with latitude and temperature as well as nutrient inputs into the system [28]. In comparison, there were no significant differences among our samples, however, GW1 showed lower richness and overall phylogenetic distance than the other samples, but similar evenness with GW2. The alpha diversity of these groundwater wells is greater than has been reported for other groundwater aquifers, with Shannon Index values below 4 and as low as 0.47 having been reported for some [29,30]. The landfill leachate well and composite leachate cistern sample diversities are consistent with the findings of Stamps *et al.* (2016), who showed high richness and evenness across the 19 U.S. landfills in their study. Similarly, the leachate richness and evenness is consistent with municipal wastewater values in Belgium (4.71<*H*<5.26 for bacteria) and China (5.80<*H*<6.23) [31,32]. The implication of these high alpha diversity values is that the landfill microbial communities consist of phylogenetically diverse microorganisms that have relatively equal abundances at the species level regardless of the presence of dominant phyla.

### Diversity of exact sequence variants

The majority (73.82%) of the ESVs are limited to one sample, suggesting that each site within the landfill hosts its own unique community. Of the 2,102 ESVs shared between at least 2 sites, 1,135 are shared between CLC_T1 and CLC_T2, suggesting that there is some proportion of the community in the composite leachate cistern that is maintained over at least a one-week interval or otherwise continually entering the cistern from the leachate wells. LW1 and LW2 consistently share more ESVs with CLC_T1 and CLC_T2 than LW3, suggesting LW1 and LW2 contribute greater amounts of leachate to the composite leachate cistern (Supplemental Table 1). Interestingly, GW2, the unimpacted well, shares more ESVs in common with the leachate wells and the composite leachate cistern than GW1, the impacted well, with 399 shared ESVs to 186 ESV, respectively (Supplemental Table 1). This is in contrast to the phylum-level differences seen for GW2 compared to landfill samples, and suggests there is some interconnectivity even between the unimpacted groundwater and the leachate – most likely caused by groundwater infiltrating the landfill. Leachate near the LW3 location is known to impact groundwater near GW1and this is supported by the higher number of shared ESVs between GW1 and LW3 than any of the other sites (Supplemental Table 1). The high number of ESVs present at only one site indicate rare operational taxonomic units, as described by Köchling *et al.* (2015) and Cardinali-Rezende *et al.* (2016), may play an oversized role in landfill microbial communities. Rare or non-prevalent organisms are hypothesized to act as seeder or starter communities during environmental changes or disturbances. Under this hypothesis, the landfill is undergoing a constant state of disturbance and change, driving the extreme heterogeneity observed.

### Microbial diversity of groundwater wells

The two groundwater wells allow a comparison between a natural groundwater environment and a leachate contaminated environment. GW1 is closer to the active landfill and leachate from the region around LW3 is leaking into the groundwater near GW1 (Figure 1). GW2 is further from the active landfill and is embedded in a region of the aquifer that shows no evidence of contamination from the landfill leachate (Figure 1, top left corner). Landfill leachate solubilizes a number of potentially harmful chemicals [11], and can be enriched with carbon and nitrogen [9]. Leachate leakage is reflected in the geochemistry data, with a number of metals and volatile compounds detected at GW1 but not GW2 (Figure 7). The higher concentration of 1,4-dioxane, vinyl chloride, and chloroethane compounds in GW1 in comparison to the leachate wells (Figure 7) may be due to loss of the landfill microorganisms capable of degrading those compounds in the aquifer near GW1. Lu *et al.* (2012) showed that when landfill leachate contaminates groundwater, the landfill microbes are unable to survive in the more dilute groundwater, and the addition of chemicals from the leachate negatively impacts the native groundwater microorganism diversity. Some natural attenuation of contaminants can occur in aquifers with leachate plumes, but more information regarding the functional abilities of these microbial communities is needed to understand the dynamics of these polluted systems [9,33].

### Functional diversity

Although the available geochemical data limit our ability to draw conclusions about the effects of geochemical parameters on the overall microbial heterogeneity of the landfill, some associations can be made for particular ESVs. Approximately 78.12% of the ESVs were classified to the genus level, either with an accepted genus name (23.33%) or other genus level grouping (54.79%). For ESVs classified as known genera, some interactions with the site geochemistry can be inferred based on current knowledge of those genera and the metabolisms utilized by their members. Three genera with either high abundance and/or prevalence that are known to interact with particular compounds are discussed below as examples of bacteria that are likely affected by the site’s geochemical heterogeneity.

### Genus *Sulfurovum*

One abundant genus in the landfill whose members have potential contaminant degradation capabilities is *Sulfurovum.* One abundant ESV, ESV_6 was classified to the *Sulfurovum* genus and is present in all samples except LW1. Four other ESVs were also classified as *Sulfurovum* at lower abundances bringing the total abundance for the genus to 1.43% (mean: 1.37%, max: 3.11% in CLC_T2). Two high quality *Sulfurovum* metagenome bins were identified: CLC_T1_32_1 and LW2_46_1. Characterized *Sulfurovum* bacteria are sulfur oxidizing bacteria and have been previously found in contaminated groundwater, with some participating in the communal anoxic degradation of benzene [34,35]. The high quality bin from LW2 encodes an ethylbenzene dehydrogenase, an enzyme used for the anaerobic degradation of hydrocarbons [36]. Benzene is present at LW3 and has historically been present at LW2 (Supplemental Table 2, Figure 7). The abundance of *Sulfurovum* at impacted or historically impacted sites and the presence of the ethylbenzene dehydrogenase gene suggests that this population may be capable of degrading benzene.

### Genus *Proteiniphilum*

*Proteiniphilum* is an abundant and diverse lineage in the landfill. Members of the *Proteiniphilum* are associated with the degradation of organic waste. ESV_5 and ESV_80 were classified to the genus *Proteiniphilum* and are present in all samples except GW1. An additional 29 ESVs classified as *Proteiniphilum* were present in various samples excepting GW1, bringing the total abundance for the genus to 2.07% (mean: 1.81%, max: 3.96% in CLC_T2). Four high-quality bins from the metagenomes were identified as *Proteiniphilum*: CLC_T2_14_7, CLC_T2_41_1, CLC_T1_27_3, and LW3_25_1. Bacteria in this genus are proteolytic [37] and the prevalence and abundance of this genus suggests a potential enrichment of organic material at these sites. Both ESV_5 and ESV_80 are present in LW2, which has higher ammonia-N and total Kjedahl nitrogen (ammonia and ammonium) (Supplemental Table 2) than all other sites. ESV_5 and ESV_80 are also present in LW1, which does not have the same concentration of nitrogen species as LW2, but which does have elevated nitrogen in comparison to the groundwater wells and LW3. This genus has also been identified at high abundance in the organic waste bioreactors studied by Cardinali-Rezende *et al.* (2016), which also had high ammonia levels. One bin CLC_T2_27_3 encodes an ATP-dependent Clp protease, an enzyme used in the degradation of proteins, while two bins, CLC_T1_27_3 and CLC_T2_14_7, encode a dinitrogenase iron-molybdenum cofactor, a critical component of the enzyme which catalyzes the reduction of dinitrogen to ammonium [38,39]. Both of these bins also encode ADP-ribosylglycohyrdolase which activates dinitrogenase reductase [40]. The presence of these components suggest that that the populations associated with these ESVs may be contributing to protein degradation and/or ammonium production in the landfill, but without evidence for complete pathways this remains hypothetical.

### Genus *Ferritrophicum*

Another genus that was relatively abundant in the landfill and has members associated with contaminant degradation is *Ferritrophicum.* Four ESVs, including ESV_4, were classified to the genus *Ferritrophicum*, which was detected at high abundance in LW3 (8.87% in LW3, 1.27% across the landfill). No metagenome bins were identified as *Ferritrophicum*. The only other recorded observation of *Ferritrophicum* species was by Weiss *et al*. (2007) who isolated the Fe(II)-oxidizing bacteria from the rhizosphere of wetland plants. There is iron present in each of the landfill leachate wells, likely in the form of Fe(II), but it is at the lowest concentration in LW3 (Supplemental Table 2). These *Ferritrophicum* bacteria may be important for iron cycling in LW3 and pose an interesting avenue for future study to understand iron cycling in the landfill and expand our knowledge of this rarely observed genus [41].

## Conclusions

The phylum level profiles for the composite leachate cistern, leachate wells, and GW1 are consistent with previous landfill microbial community studies, with *Bacteroidota, Firmicutes*, and *Proteobacteria* among the most abundant phyla. Using metagenomic sequencing, we additionally identified *Patescibacteria* as one of the dominant phyla in the landfill, a group that may have been missed in previous studies relying on 16S rRNA gene amplicon sequencing. At the species/ESV level, microbial heterogeneity is markedly higher than previously reported for landfill environments, with little overlap between communities separated by short distances or one week in time. Geochemical conditions showed high variance across the site, and were generally uncorrelated with microbial community memberships. Taken together, our findings have implications for waste management strategies, including targeted remediation efforts, as establishing populations and activities of interest will be challenging given the dynamic nature of the landfill microbial communities.

## Materials and Methods

### Sample collection

In the initial sampling event on July 14, 2016, a sample was collected from the composite leachate cistern by filtering the leachate through a 0.2 μm poly-ethersulfone filter followed by a 0.1 μm poly-ethersulfone filter in series (CLC_T1_0.2 and CLC_T1_0.1, respectively). Both filters were kept for DNA extractions. On July 20, 2016, a larger-scale sampling was conducted, sampling the composite leachate cistern (CLC_T2), three leachate wells (LW1, LW2, LW3), and two groundwater wells (GW1, GW2). Leachate and groundwater samples were collected by pumping liquid through a filter apparatus with a 3 μm glass fiber pre-filter in series with a 0.1 μm poly-ethersulfone filter until filters clogged. The pre-filter was discarded while the 0.1 μm filters carrying the microbial biomass were kept. All filters were frozen on dry ice in the field and transferred to a −80 °C freezer until processed. DNA was extracted from the biomass using the Powersoil DNA extraction kit (MoBio) following the manufacturer’s instructions with one modification: filters were sliced into pieces and added to the bead tube in place of a soil sample.

Relevant measurements for volatile and non-volatile compound concentrations at the leachate and groundwater wells are conducted each year in October and April by a contracted consulting company. For 2016, the average values for these two sampling points were used to estimate compound concentrations in July, the time of microbial biomass sampling (Supplementary Table 1). The impacted groundwater well, GW1, did not have current non-volatile concentration measurements available. For this well, measurements from 2011 were included for comparison purposes only (Supplementary Table 1). No geochemical measurements were available for the composite leachate cistern.

### Sequencing

All eight samples were sent to the JGI for 16S rRNA gene amplicon sequencing: LW1, LW2, LW3, CLC_T1 0.1 μm and 0.2 μm filters, CLC_T2, GW1, and GW2. The JGI amplified the V4 region of the 16S rRNA gene using the forward primer 515FB (5’-GTGYCAGCMGCCGCGGTAA-3’) and the reverse primer 806RB (5’-GGACTACNVGGGTWTCTAA-3’) using in-house protocols. Amplicons were sequenced on the MiSeq platform (Illumina) with all appropriate negative and positive controls, and reads were quality control checked using the iTagger pipeline [42].

Six DNA samples were sent to the US Department of Energy’s Joint Genome Institute (JGI) for metagenomic sequencing, assembly, and annotation: LW1, LW2, LW3, CLC_T1 (0.2 μm filter), CLC_T2, and GW1. The CLC_T1 and GW2 0.1 μm filters resulted in insufficient DNA and were not sent for metagenomic sequencing. The JGI sequenced the metagenomes using the HiSeq platform (Illumina). Metagenomes were annotated using the DOE-JGI Metagenome Annotation Pipeline (MAP v.4) [43].

### Phylogenetic trees

All assembled and annotated 16S rRNA genes in the landfill metagenomes were downloaded from the JGI IMG server (IMG Genome IDs: CLC_T1: 3300014203, CLC_T2: 3300014206, LW1: 3300014204, LW2: 3300015214, LW3: 3300014205, GW1: 3300014208). Genes were sorted by length in Geneious 11.0.5 (https://www.geneious.com) and curated to a minimum length of 600 bp. The landfill metagenome-derived 16S rRNA genes as well as a reference set of 16S rRNA genes from known organisms to the SILVA SINA algorithm [44] for alignment. Unaligned bases at the ends of the genes were removed and sequences below 70% identity to a reference sequence were automatically removed from the dataset by SINA. To curate the SINA alignment, columns containing 97% or more gaps were removed, a region of poor alignment was manually trimmed from the 3’ end, and sequences falling below 600 bp post-trimming were removed. A phylogenetic tree was inferred using FastTree in Geneious to check for poorly aligned or divergent sequences. In this processing, 195 sequences were removed that did not meet quality standards. The final 16S rRNA gene alignment included 1,903 reference sequences and 2,306 sequences from the metagenome samples, and had 1,521 positions. A high-quality phylogenetic tree was inferred from the curated final alignment using RAxML-HPC2 8.2.12 [45] on Cipres [46] under model GTRCAT, with 100 alternative bootstrap iterations run from 100 starting trees.

All amino acid sequences for 16 syntenic, universally-present, single copy ribosomal protein genes (RpL2, L3, L4, L5, L6, L14, L15, L16, L18, L22, L24 and RpS3, S8, S10, S17, S19) for the landfill metagenomes were downloaded from the JGI IMG server using annotation keyword-based identification [47]. Ribosomal protein datasets were screened for the Archaeal/Eukaryotic type, which were removed, as were short (<45 aa) sequences. Each individual protein set was aligned with a reference set of genes [48] using MAFFT 7.402 [49] on Cipres. Alignment columns containing ≥95% gaps were removed using Geneious. IMG-derived sequence names were trimmed to 8 digits after the metagenome code (e.g. Ga0172377_100004578 → Ga0172377_10000457) to remove gene-specific identifiers and allow for concatenation by scaffold name. The protein gene alignments were concatenated in numeric order (L2 → L24, followed by S3 → S19). Concatenated sequences that contained less than 50% of the total expected number of aligned amino acids were removed. The final alignment was 3,452 columns long and contained 2,914 reference organisms and 1,265 scaffolds from the metagenome samples. A phylogenetic tree was inferred using RAxML-HPC Blackbox on Cipres using the following parameters: sequence type - protein; protein substitution matrix – LG; and estimate proportion of invariable sites (GTRGAMMA + I) – yes [45,46].

### 16S rRNA amplicon sequence analyses

The demultiplexed and barcode-trimmed 16S rRNA gene amplicons from the JGI were analyzed using QIIME2 [50]. Forward and reverse reads were separated using khmer [51]. Primers were trimmed from the forward and reverse reads using cutadapt in QIIME2 [52]. The forward reads were truncated at 231 base pairs and the reverse reads at 230 base pairs based on the quality score visualization produced by QIIME2 in the demux summary step. Reads were denoised using paired denoising in DADA2 within the QIIME2 platform which also merges the reads [53]. Sequence variants were determined using DADA2 and summarized using feature-table summarize in QIIME2. Taxonomic assignment of the 16S rRNA gene amplicons was based on a phylogenetic tree produced by QIIME2 in which the taxonomy classifier was trained with the SILVA 99% taxonomy classification for the 16S rRNA gene from the April 2018 SILVA 132 release [54]. Phylum names were updated as per the GTDB database taxonomy changes by Parks *et al.* (2018) for diversity comparisons.

All scaffolds >2,500 bp were included in the binning process. The binning algorithm CONCOCT [55] was used in Anvi’o [56] to automatically cluster each metagenome’s scaffolds using a combination of scaffold tetranucleotide frequencies and read-mapped coverage data from all six metagenomes. Gene annotations were imported from the JGI annotations, overriding the automated annotation pipeline in Anvi’o. The bins were manually refined for the six metagenomes using Anvi’o, focusing on completion and quality metrics to guide bin refinements. High quality bins were considered those with greater than 70% completion and less than 10% redundancy.

### Diversity analyses

Diversity analyses on the 16S rRNA gene amplicon exact sequence variants (ESVs) identified by QIIME2 [50] included the alpha diversity metrics Faith’s phylogenetic diversity [57] and Pielou’s evenness [58], calculated based on four sample types: impacted groundwater well, unimpacted groundwater well, leachate well, and composite leachate cistern. A Shannon diversity index analysis with rarified sequence depth of 53,518 was conducted using QIIME2 and visualized using phyloseq [22] in R. A Chao1 statistic was not calculated, as data processing with QIIME2 and DADA2 removes all singleton ESVs, which the Chao1 statistic requires. For beta diversity measures, full ESV and taxonomy tables were input to unweighted and weighted UniFrac distances principle coordinate analyses, calculated using phyloseq and visualized in R for all samples. The prevalence across samples of ESVs with a count of 2 or more and belonging to phyla with relative abundance greater than 1% or present in multiple sites was determined using phyloseq and visualized in R. The phyla with ESVs present in five or more sites was visualized using ggplot2 in R.

A principal component analysis (PCA) was conducted using vegan [59] in R for all 16S rRNA gene amplicon ESVs and the16S rRNA gene amplicon ESVs present at five or more sites. The ESV count data was Hellinger transformed to reduce the weight of ESVs with low counts and zeros. The leachate wells and the two groundwater well samples were included in the PCA to allow for comparison with environmental parameters, which are available for those sites. Environmental data was not available for the composite leachate cistern site and so CLC samples were included in the analysis only for comparison with the other samples.

Metagenome-derived sequences were classified at the phylum level based on their placement within reference clades on the 16S rRNA and concatenated ribosomal protein phylogenetic trees. Metagenome sequences placing outside of or between phyla were assigned to either “Unclassified Archaea” or “Unclassified Bacteria” as appropriate. Phylum names were updated from the NCBI taxonomy to conform to the GTDB database taxonomy by Parks *et al.* (2018). Bins were identified at the phylum level using the scaffold assignments from the 16S rRNA gene and concatenated ribosomal protein phylogenetic trees. Bin abundances were determined using the average fold coverage data for all scaffolds in the bin. Phylum abundance per sample was calculated by summing the average fold coverage data for each scaffold on the tree assigned to the phylum, where the scaffold acts as a proxy for the underlying microbial population. Microbial diversity comparisons were visualized using stacked bar plots produced using ggplot2 in R [23].

### Chemical data analyses

Chemical measurements provided by the Southern Ontario landfill 2016 annual report were used to determine variance of non-volatile and volatile compounds over time for the three leachate wells and the unimpacted groundwater well. GW1 only has non-volatile compound measurements for one time point in 2011 and so variance could not be calculated. Non-detects, where a compound, if present, is below the detection limit, were treated as zeros. The measurements were log transformed and visualized in a heatmap using heatmap3 [60] in R. Metal and volatile compounds detected in a majority of samples were used for further analysis. The measurements from April and October of 2016 were averaged to estimate the concentrations at the time of microbial biomass sampling.

PCA for the metals and volatile compounds were conducted using vegan [59] in R. The metal and volatile compound concentrations were square root transformed to reduce the range of the values as different compounds differed in concentration by orders of magnitude (Supplemental Table 2). Data for leachate wells and the two groundwater well samples were included for the volatile analysis, but GW1 was excluded from the non-volatile compound analysis as no data were available for that site in 2016. A PCA was also conducted using vegan in R for the other geochemical parameters measured at the sites that are not characterized as non-volatile or volatile compounds (*e.g.,* total dissolved solids (TDS)).

## Supporting information

Supplemental File 1 - trees

## Figure Legends

**Supplemental Figure 1: Alpha diversity analyses by sample type for Faith’s Phylogenetic Diversity and Pielou’s Evenness.** A) Faith’s phylogenetic diversity. A Kruskal-Wallis test for all groups is not significant (H=2.77 and p=0.43). Kruskal-Wallis pairwise tests are also not significant for all pairs (p≥0.18). B) Pielou’s evenness. All sample types show high levels of evenness among their respective species (J’ > 0.74). A Kruskal-Wallis test for all groups is not significant (H=3.22 and p=0.36). Kruskal-Wallis pairwise tests are also not significant for all pairs (p≥0.18).

**Supplemental Figure 2: Phyla with 16S rRNA gene amplicon ESVs in five or more sites.** The total count of ESVs present in five or more sites was summed by phylum.

**Supplemental File 1: Newick trees for concatenated ribosomal protein tree and 16S rRNA tree used to identify metagenome-derived microbial community composition.**

## Declarations

### Ethics approval and consent to participate

There were no ethics approvals or consents required for the research presented here.

### Consent for publication

All relevant parties have confirmed consent for publication, including all co-authors.

### Availability of Data and Material

The assembled and annotated Southern Ontario metagenomes are deposited on IMG with the following IMG Genome IDs (Taxon Object IDs): 3300014203 (CLC1_T1), 3300014206 (CLC1_T2), 3300014204 (LW1), 3300015214 (LW2), 3300014205 (LW3), and 3300014208 (GW1).

The 16S rRNA amplicon sequences have been deposited in the NCBI SRA archive under the bioproject PRJNA706007 and Biosamples SAMN18111220-7.

### Authors’ contributions

AS and LAH conceived of the study. AS and LAH conducted fieldwork and generated samples. AS analyzed the data and wrote the first draft of the manuscript. AS and LAH edited the manuscript. LAH secured funding to support the research.

## Acknowledgements

We thank members of the Hug Research Group, past and present, for help with landfill sampling as well as metagenome binning, which was done collectively. We are grateful to the municipality that allowed access to the landfill for sampling including the contracted consulting company and the summer intern who accompanied us on our sampling trips, anonymity requested.

