## Supplemental methods and figures for "High diversity and heterogeneity define microbial communities across an active municipal landfill"

### Supplemental Tables and Figures

**Supplemental Table 1: Number of shared 16S rRNA gene amplicon ESVs between samples.**

| | CLC_T1 | CLC_T2 | LW1 | LW2 | LW3 | GW1 | GW2 |
|---|---|---|---|---|---|---|---|
| CLC_T1 | | 1135 | 384 | 301 | 223 | 27 | 280 |
| CLC_T2 | 1135 | | 431 | 486 | 234 | 13 | 295 |
| LW1 | 384 | 431 | | 195 | 476 | 59 | 132 |
| LW2 | 301 | 486 | 195 | | 119 | 8 | 142 |
| LW3 | 223 | 234 | 476 | 119 | | 131 | 116 |
| GW1 | 27 | 13 | 59 | 8 | 131 | | 46 |
| GW2 | 280 | 295 | 132 | 142 | 116 | 46 | |

**Supplemental Table 2: Averaged environmental variable concentrations based on measurements from April and October 2016 for the groundwater wells and the three leachate wells.**

| *Environmental Parameter* | *units* | *GW1** | *GW2* | *LW1* | *LW2* | *LW3* |
|---|---|---|---|---|---|---|
| Arsenic | mg/L | 0.02 | 0 | 0 | 0.04 | 0 |
| Boron | mg/L | 0.79 | 0.01 | 2.10 | 7.53 | 1.45 |
| Cadmium | mg/L | 0 | 0 | $0.10e^{-3}$ | 0 | 0 |
| Calcium | mg/L | 240 | 86.50 | 175 | 93.25 | 155 |
| Chromium | mg/L | 0 | 0 | 0.04 | 0.40 | 0.02 |
| Copper | mg/L | 0 | 0 | 0.01 | 0.36 | 0.03 |
| Iron | mg/L | 40 | 0.37 | 56.50 | 11.78 | 10.55 |
| Lead | mg/L | 0 | 0 | 0.01 | 0.05 | 0 |
| Magnesium | mg/L | 38 | 24 | 145.50 | 145 | 58.50 |
| Manganese | mg/L | 1.20 | 0.03 | 0.32 | 0.29 | 0.67 |
| Mercury | mg/L | 0 | 0 | 0 | $0.30e^{-3}$ | 0 |
| Nickel | mg/L | 0.01 | 0 | 0.05 | 0.36 | 0.02 |
| Potassium | mg/L | 16 | 1.40 | 345 | 917.50 | 79.50 |
| Sodium | mg/L | 76 | 4.55 | 615 | 2525 | 200 |
| Zinc | mg/L | 0 | 0 | 0.06 | 0.28 | 0.01 |
| Alkalinity total (as $CaCO_3$) | mg/L | 687 | 250 | 2800 | 7025 | 1200 |
| Ammonia-N | mg/L | 18 | 0 | 275 | 1375 | 123.50 |
| Chloride | mg/L | 180 | 6.70 | 655 | 3050 | 255 |
| Nitrite | mg/L | 0 | 0 | 0.01 | 46.24 | 0.01 |
| Sulfate | mg/L | 17 | 60.50 | 0 | 70 | 51.50 |
| Total dissolved solids | mg/L | 1040 | 315 | 3255 | 9657.50 | 1345 |
| Total Kjeldahl nitrogen | mg/L | 20 | 0.11 | 315 | 1925 | 135 |
| Un-ionized ammonia | mg/L | 0.03 | 0 | 2.32 | 99.88 | 0 |
| 1,4-Dichlorobenzene | ug/L | 0 | 0 | 0 | 0 | 11.50 |
| Benzene | ug/L | 0.69 | 0 | 13.50 | 0 | 5.50 |
| Chlorobenzene | ug/L | 0 | 0 | 6 | 0 | 3.80 |
| Ethylbenzene | ug/L | 0.13 | 0 | 275 | 8.25 | 17.50 |
| m&p-Xylenes | ug/L | 0.07 | 0.06 | 1040 | 26.25 | 200 |
| o-Xylene | ug/L | 0.06 | 0 | 250 | 17.05 | 11.20 |
| Toluene | ug/L | 0 | 0 | 17 | 0 | 0 |

*All measurements for GW1 except the volatile compounds (1,4-dichlorobenze to toluene) are from April 2011 and are for comparison only. These values were not included in the PCA analysis.

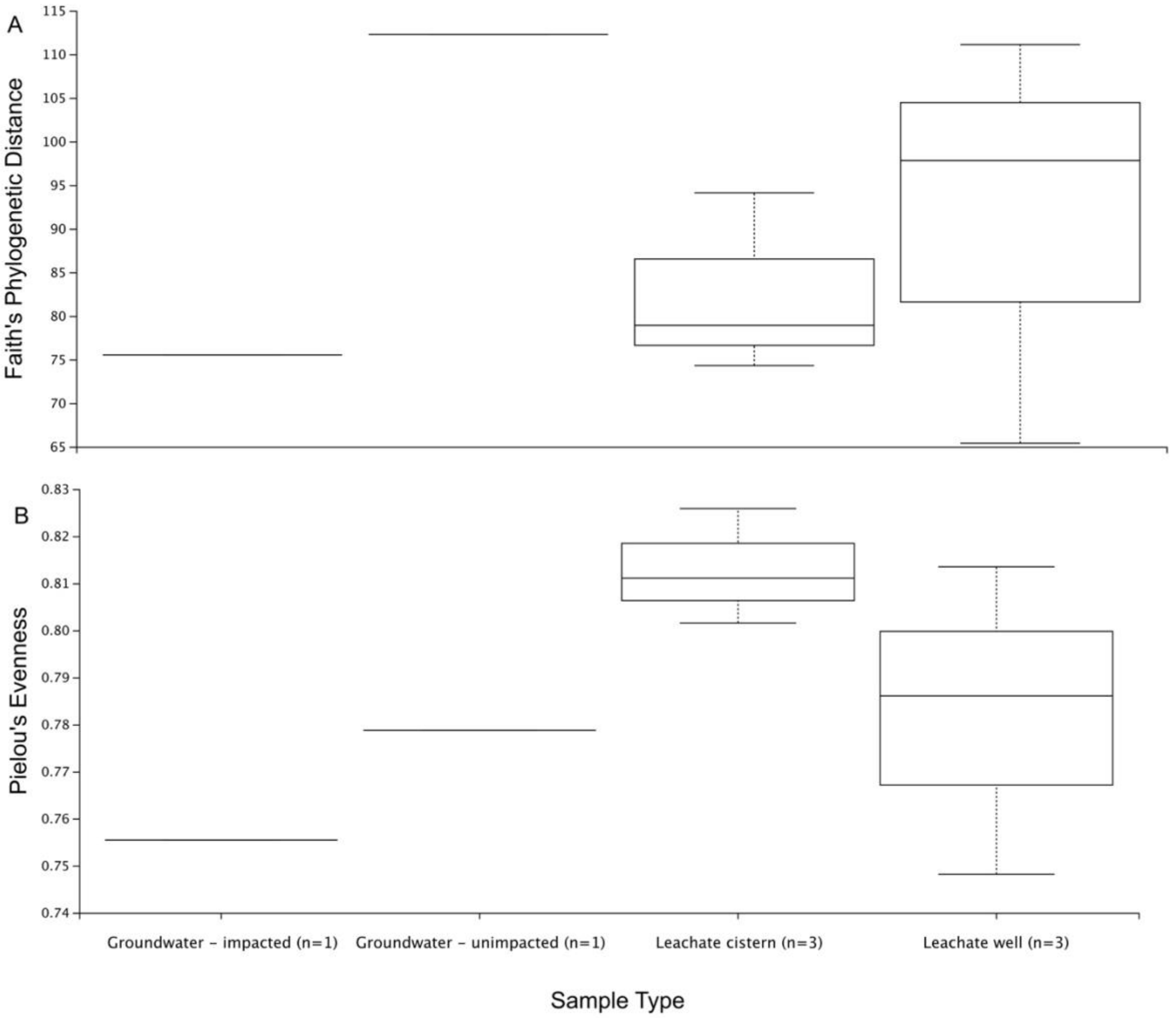


**Supplemental Figure 1: Alpha diversity analyses by sample type for Faith's Phylogenetic Diversity and Pielou's Evenness.** A) Faith's phylogenetic diversity. A Kruskal-Wallis test for all groups is not significant ($H=2.77$ and $p=0.43$). Kruskal-Wallis pairwise tests are also not significant for all pairs ($p \geq 0.18$). B) Pielou's evenness. All sample types show high levels of evenness among their respective species ($J' > 0.74$). A Kruskal-Wallis test for all groups is not significant ($H=3.22$ and $p=0.36$). Kruskal-Wallis pairwise tests are also not significant for all pairs ($p \geq 0.18$).

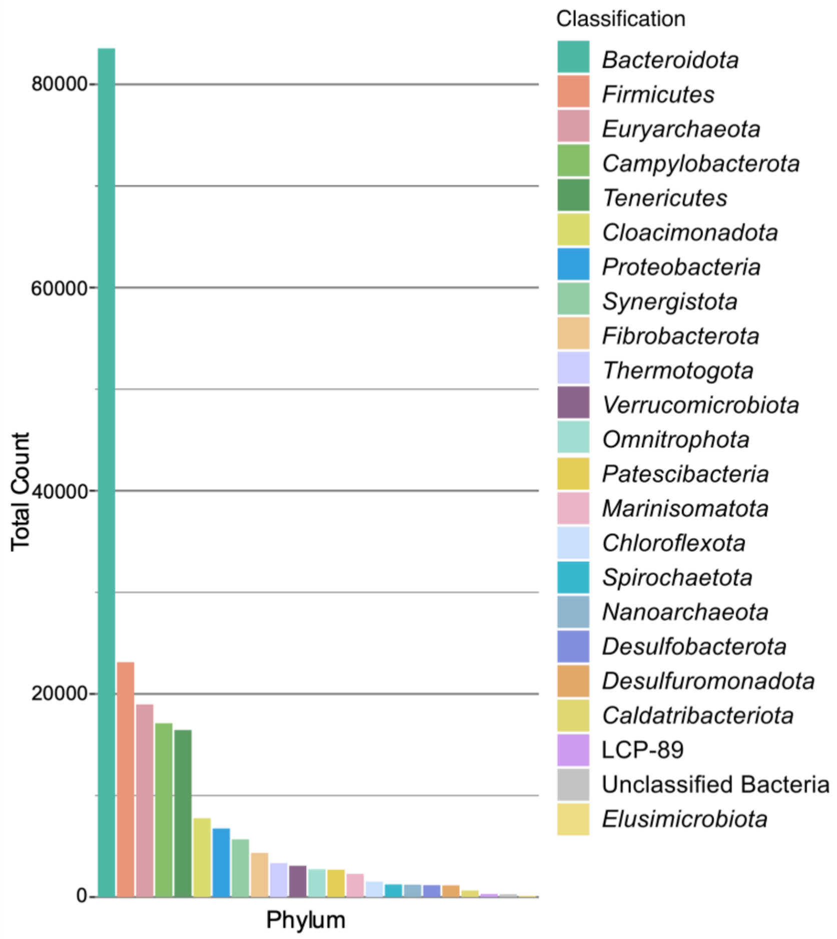


**Supplemental Figure 2: Phyla with 16S rRNA gene amplicon ESVs in five or more sites.** The total count of ESVs present in five or more sites was summed by phylum.
